# Functional analysis of Scratch2 domains: implications in the evolution of Snail transcriptional repressors

**DOI:** 10.1101/410761

**Authors:** Tatiane Yumi Nakamura Kanno, Mariana Soares Fogo, Carolina Purcell Goes, Felipe M. Vieceli, Chao Yun Irene Yan

## Abstract

The Snail superfamily of transcription factors have a modular organization and their similarities and divergences are the basis for subdividing the superfamily into the Snail1/2 and Scratch families. As it is generally accepted that the Snail and Scratch families originated through gene duplication, understanding the functional contribution of each module could provide us with further insight about the molecular and functional evolution of the Snail superfamily. Thus, in this work, we investigated the function of the SNAG and SCRATCH domains in chicken Scratch2. Through evolutionary comparison analysis we identified a novel HINGE domain that lies between the SNAG and SCRATCH domain. Similar to members of the Snail1/2 families, Scratch2- mediated transcriptional repression requires SNAG and nuclear localization requires the zinc-finger domain. We also identified a novel HINGE domain that lies between the SNAG and SCRATCH domain. HINGE is highly conserved in amniotes. Single mutations of the conserved Tyrosine and Serine residues of HINGE downregulated Scratch2-mediated transcriptional repression. This effect depended on the presence of the SCRATCH domain.

## Introduction

The members of the Snail superfamily of transcription factors are key players in multiple embryological and pathological events (reviewed in [1] acting mainly as transcriptional repressors [2–4]. Similarities and divergences in their sequences subdivides the superfamily into the Snail1/2 and Scratch families. This subdivision was originally based on sequence homology comparison, but it clearly correlates with functional differences as well. Both Snail and Scratch families modulate cell adhesion and migration. Whilst the effect of Snail is broader [5–7], Scratch is limited to neural development, regulating neuronal migration during cortical formation, promoting neural fate and repressing neural precursor cell death [8–10].

All Snail superfamily members have a modular organization, including a conserved carboxy-terminal region containing zinc-fingers and a SNAG (<u>Sna</u>il/<u>G</u>fi-1) domain in the amino-terminus. Scratch and Snail2 proteins present distinct conserved domains between the SNAG and zinc-fingers domains, called SCRATCH and SLUG domains, respectively. The correlation between domains and function has been much better characterized in the Snail than in the Scratch family. The zinc-finger region of Snail proteins recognizes a consensus DNA motif containing a core of six nucleotides known as E-boxes [11], and the SNAG domain of Snail1 interacts with a histone lysine-specific demethylase, repressing target genes [12]. The SLUG domain has also been reported to contribute to repression [13] and members of the Snail family that lack a SNAG domain repress through an alternative domain known as the CtBP domain [14].

In Scratch1, the repressor domain has also been attributed to the amino-terminus of the protein, but deletion of the SNAG domain in human Scratch1 does not reduce its ability to repress E-box-driven transcription [4]. Further, the Scratch family lacks an obvious alternative module such as the CtBP domain. Thus, the repressor domain in the Scratch family remains undefined. As it is generally accepted that the Snail and Scratch families originated through gene duplication, with the signature domains of each family arising through divergence of an ancestral gene [15], understanding the functional contribution of each module could provide us with further insight about the molecular and functional evolution of this superfamily.

In this work, we have investigated the biological role of different domains in Scratch2 (Scrt2) through deletion and point mutations in the chicken orthologue. Our data suggest that Scrt2 requires SNAG for its repression activity. Also, we identified another conserved motif-HINGE-that co-evolved with the SCRATCH domain. Scrt2-mediated transcriptional repression can be modulated by the SCRATCH domain through modifications in the HINGE -a novel region conserved in the Scrt2 subfamily.

## Material and methods

### Generation of mutated and truncated proteins

The pCIG-MYC-*cScrt2* containing chicken Scrt2 cDNA (JN982016.1) was previously cloned in our laboratory [16] and has been used as a template for the generation of wild-type truncated constructs. Full-length N-terminal FLAG-tagged single mutation constructs harboring substitutions tyrosine 77 to phenylalanine (*cScrt2-*Y77F) or glutamate (*cScrt2-*Y77E), or serine 78 to alanine (*cScrt2-*S78A) or aspartate (*cScrt2-*S78D) were synthesized by GenScript USA Inc. (Piscataway, NJ). N-terminal FLAG-tagged *cScrt2*ΔSCRATCH (aa 97-116 deletion), the full-length double mutation constructs containing both Y77F and S78A (*cScrt2-*YS/FA) or Y77E and S78D (*cScrt2-*YS/ED), and Y77 or S78 substitutions in the absence of the SCRATCH domain (YFΔSCRATCH or SAΔSCRATCH) were synthesized by Integrated DNA technologies (IDT). All commercially purchased constructs were subcloned into pCIG, where expression of a bicistronic RNA with nuclear GFP reporter is driven by chicken beta-actin promoter [17]. The sequences for *cScrt2*ΔZnF (aa 1-127), *cScrt2*ΔN (aa 124-276), *cScrt2*ΔSNAG (aa 10-276), *cScrt2*S78AΔSNAG (aa 10-276, S78A) and *cScrt2*YS/FAΔSNAG (aa 10-276, Y77F and S78A) were PCR amplified flanked by EcoRI and SmaI sites and subcloned into pCIG or pMES. The latter differs from pCIG in that it the GFP lacks a nuclear localization signal and remains in the cytoplasm [18].

### In ovo electroporation

Chicken embryos at stage HH10-HH12 [19] were electroporated with *cScrt2*WT, *cScrt2*- Y77F and *cScrt2*-S78A. Electroporated cells were identified by the presence of GFP. Briefly, a small window was made at the top of the egg shell to reach the embryo. The embryos were visualized with sterile Indian ink 10% (diluted in Howard Ringer’s saline solution) injected under the blastoderm. The plasmid solution (concentration of 3 μg/ml) containing the inert tracer FastGreen 0.2% was injected into the truncal neural tube lumen. Then, the platinum electrodes were placed at a distance of 4 mm flanking the neural tube and 5 pulses of 20 V with 30 ms of length and 100 ms of interval were administered [17,20]. Embryos were re-incubated and collected 24 hours later.

### HEK293T culture and transfection

Established HEK293T cells were cultured in DMEM supplemented with 10% FBS and 1% antibiotics (streptomycin 5 μg/ml and penicillin 5 U/ml). Transfection was performed with 2 ug Lipofectamine 2000 (Invitrogen) and 0.8 μg of the DNA construct per well in 24-well plates for 4 hours in Opti-MEM medium without antibiotics. After 4 hours of transfection, the cells were washed with serum-free medium and fed with complete DMEM medium. The cells recovered for 16–18 hours before fixation.

### Immunofluorescence

Embryos were fixed in PBS/paraformaldehyde 4% for 30 minutes and cryoprotected with 20% sucrose overnight at 4°C and embedded in an OCT-20% sucrose mixture (1:1) prior to sectioning in cryostat at 10 μm. We sectioned the trunk region of the embryo between the limb buds. The slides were dried for 30 minutes at 37°C, fixed in PBS/paraformaldehyde 4% for 20 minutes, washed three times of 10 minutes with PBS and blocked for 1 hour with 3% NGS and 1% BSA diluted in PBST (PBS containing 0,1% Triton X-100), followed by incubation with antibodies. Coverslips containing transfected HEK293T cells were washed once with PBS for 10 minutes and then fixed in PBS/paraformaldehyde 4% for 20 minutes. Next, the coverslips were washed with PBS and incubated with blocking solution (PBS containing 0,1% Triton X-100 and 3% NGS), followed by incubation with antibodies. Primary antibodies were diluted in the block solution and applied on sections or cells overnight at room temperature in a humidified chamber. In the next day, the slides or coverslips were washed with PBS and then incubated with the secondary antibody for 2 hours at room temperature. DAPI was added to the secondary antibody solution for nuclear staining. Primary antibodies used were: anti-MYC (0,004 mg/ml – 9E10, Life Technologies), anti-GFP (0.002 mg/ml - A-11122, Life Technologies), anti-FLAG (1:250 - F3165, Sigma). Secondary antibodies used were goat anti-rabbit IgG Alexa Fluor 488 (1:500, Molecular Probes) and anti-mouse IgG Alexa Fluor 568 (1:500, Molecular Probes) or 647 (1:500, Molecular Probes).

### Luciferase assay

For these experiments, we inserted four E-Box sequences (CAACAGGTG) in *tandem* into pGL3Luc vector (Promega), generating the plasmid-test pGL3-4xE-box. This plasmid is used to indirectly measure the transcriptional activity of Scrt2 through the activity of luciferase. To perform the assay, HEK293T cells were dissociated and plated in 24-well plates at the concentration of 1.25×10^5^ cells/well. The cells were transiently co-transfected with each plasmid containing the tested constructs together with pGL3-4xE-box and pRL encoding the renilla luciferase. Renilla luciferase is transcribed independently of Scrt2 and served as a normalization factor for the assay. Control conditions were the same, except that the tested construct (pMES or pCIG) did not contain cScrt2 or its variants.

The co-transfection was performed with Lipofectamine 2000 (Invitrogen) at a final concentration of 2 μg, 0.4 μg of tested plasmids, 0.01 μg of pRL and 0,4 μg pGL3-4xE-box for 4 hours in Opti-MEM medium. After 4 hours, the medium containing the transfection solution was removed, the cells were washed with serum-free medium and cultured with complete medium. The cells were re-incubated for 16-18 hours before collection and luciferase signal was measured following the kit manufacturer’s instructions (Dual luciferase assay reporter system, Promega). Statistical analysis was performed using one-way ANOVA. The level of significance adopted was p <0.05.

## Results

### The zinc-finger domain of Scrt2 determines subcellular localization

The main activity reported for members of the Snail superfamily is transcriptional repression. This is due to the joint effect of the zinc-finger domain, mediating nuclear translocation and DNA-binding activity, and the SNAG domain, mediating the repressor activity [12]. To investigate the role of the zinc-finger domain in Scrt2 nuclear localization, we expressed a series of truncated chicken Scrt2 proteins that included or excluded the zinc-fingers, as well as the conserved domains SNAG and SCRATCH, in different combinations (Fig. 1A).

**Figure 1.**
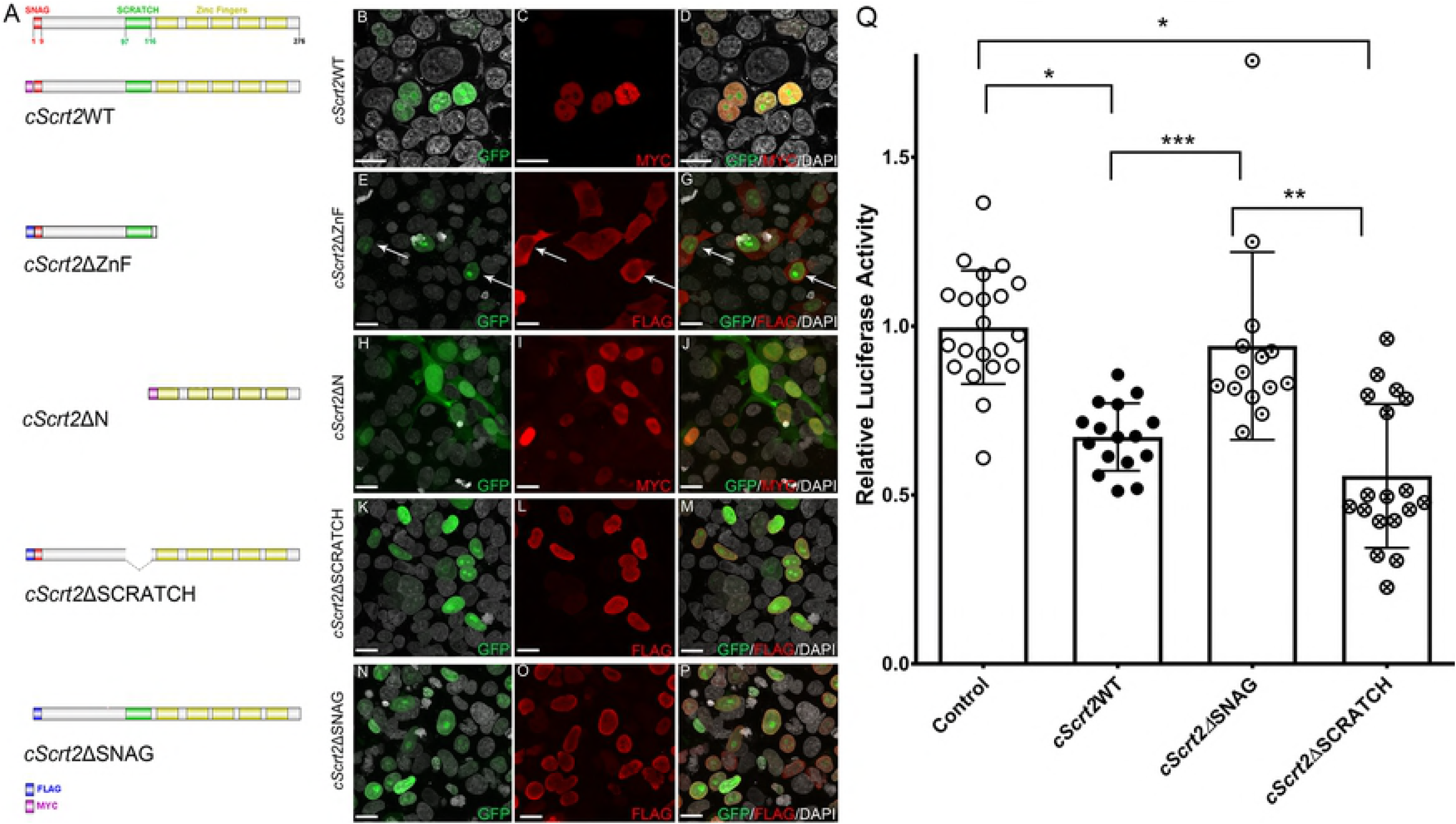
Nuclear localization of chicken Scrt2 depends on the zinc-finger domain while the transcriptional repression activity is regulated by SNAG domain. (A) Diagram on the left represents chicken Scrt2 full length protein (amino acids 1-276). The different domains are represented with a color code: the SNAG domain (1 to 9 aminoacids) in red, the SCRATCH domain (97 to 116 aminoacids) in green, and the five zinc fingers, comprising the DNA binding domain, in yellow. MYC-tag and FLAG-tag are represented in magenta and blue, respectively. *cScrt2*WT corresponds to the full sequence, with a MYC-tag at the N-terminal region. Below *cScrt2*WT are represented, in order, the diagrams for *cScrt2*ΔZnF (amino acids 1 to 127), *cScrt2*ΔN (amino acids 124 to 276), *cScrt2*ΔSCRATCH (deletion of amino acids 97-116) and *cScrt2*ΔSNAG (amino acids 10 to 276). Co-localization with nuclear GFP and DAPI indicates that expression of *cScrt2*WT (B-D), *cScrt2*ΔN (H-J), *cScrt2*ΔSCRATCH (K-M), and *cScrt2*ΔSNAG (N-P) localize to the nucleus in HEK293T. In contrast, *cScrt2*ΔZnF expression is restricted to the cytoplasm (E-G, arrows). (B) Whereas *cScrt2*WT reduced transcription factor-activated luciferase activity relative to control conditions, removal of SNAG (*cScrt2*ΔSNAG) induced luciferase activity to levels similar to control, suggesting that the absence of SNAG decreases chicken Scrt2-mediated transcriptional repression. In contrast, chicken Scrt2 lacking the SCRATCH domain (*cScrt2*ΔSCRATCH) repressed transcription with the same efficiency as *cScrt2*WT. In control conditions, HEK293T cells were transfected with pGL3-4xE-box and empty pCIG vector. *cScrt2*WT, *cScrt2*ΔZnF, *cScrt2*ΔSCRATCH and *cScrt2*ΔSNAG are inserted into pCIG vector while *cScrt2*ΔN is inserted into pMES vector. Results show the scatter plot distribution of at least triplicate samples from three different experiments, +/- standard deviation. The data was analyzed by one-way ANOVA. *p<0.0001 **p=0.0007 ***p=0.02. Scale bar: 20μm

*cScrt2*ΔZnF, lacking the zinc-finger but including the SNAG and SCRATCH domains, was found only in the cytoplasm (Fig. 1E-G, arrows) of transfected HEK293T. In contrast, all the truncations that included the zinc-finger domain, that is, Scrt2 lacking either the SNAG (*cScrt2*ΔSNAG) or SCRATCH (*cScrt2*ΔSCRATCH) domains or containing only the zinc-finger motif (*cScrt*ΔN), segregated to the nuclei of HEK293T cells (Fig. 1H-P).

As reliable Scrt2-reactive antibodies are lacking, we used epitope-tagged Scrt2 for these experiments, first assessing if the presence of the epitope tag affected Scrt2 subcellular localization. Consistent with its reported role as a DNA-binding transcription factor, MYC-tagged Scrt2 co-localized with the nuclear DAPI stain in HEK293T cells and in embryonic neural cells (Fig. 1B-D and Fig. S1).

### Scrt2 repressor activity requires the conserved SNAG domain but not the SCRATCH domain

In most members of the Snail superfamily, transcriptional repressor activity requires the conserved amino-terminus SNAG domain (Fig. 2) [13]. Accordingly, removing the SNAG domain (*cScrt2*ΔSNAG) decreased Scrt2-mediated transcriptional repression significantly (Fig. 1Q), without affecting its nuclear localization (Fig. 1O).

**Figure 2.**
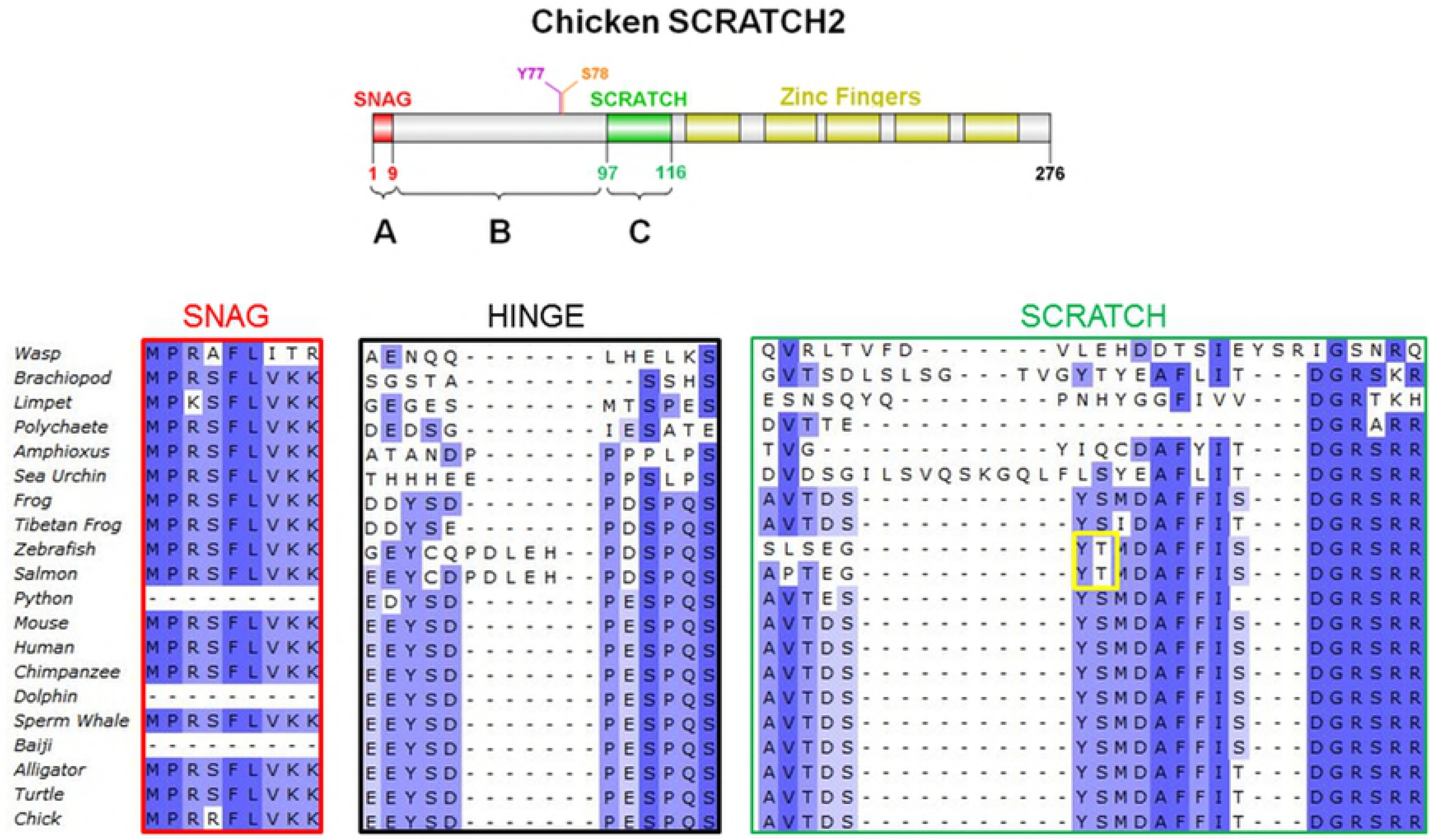
Chicken Scrt2 has conserved amino acid domains. Top diagram represents full length chicken Scrt2 with its protein domains. Chicken Scrt2 sequence conservation was analyzed in a variety of organisms with Ugene and ClustalW programs. Deeper hues reflect higher degree of conservation. SNAG (red box), HINGE (black box) and SCRATCH (green box) domains display high degree of conservation. The HINGE domain displays a strongly conserved sequence (EEYSDPESPQS, amino acids 75-85 – black box) before the SCRATCH domain, suggesting that this region might be important for chicken Scrt2 function. Accession numbers for the sequences used are in Fig. S2B. Python and Baiji sequences are partial.

Besides the SNAG and zinc-finger domains, additional domains are conserved in different branches of the Snail superfamily and are used for their phylogenetic classification. In particular, the majority of the members of the ScratchB [21] branch of the Scratch family have the SCRATCH domain. Here, we consider the SCRATCH domain as the full AVTDSYSMDAFFITDGRSRR sequence (aminoacids 97-116 in chicken Scrt2, Fig. 2). This domain lies between the SNAG and zinc-fingers domains, and its function remains unknown [21]. As SCRATCH was not required for Scrt2 nuclear localization (Fig. 1), we next evaluated the effect of removal of SCRATCH domain on transcriptional repressor activity. The truncated form *cScrt2*ΔSCRATCH displayed transcriptional repression similar to the native form *cScrt2*WT, suggesting that SCRATCH is required neither for nuclear localization nor for repressor activity (Fig. 1Q).

### HINGE domain Ser and Tyr residues are required for Scrt2 repressor activity

To further investigate the role of the SCRATCH domain, we analyzed its evolution in the context of Scrt2 proteins through sequence alignment (Fig. 2). We observed that the full SCRATCH domain co-evolved in vertebrates together with another conserved domain that we named HINGE (amino acids 75-85 in chicken Scrt2; Fig. S2A). HINGE is extremely well conserved in amniotes and contains an initial acidic-rich motif EEYSD. The acidic residues of the motif can vary between glutamate and aspartate, but the core residues Tyrosine_77_ (Y77) and Serine_78_ (S78) are maintained in most vertebrates – except fish. Furthermore, these two residues are potentially recognized by a variety of kinases (Fig. S3). As changes in phosphorylation levels modulate protein stability and repressor activity of Snail1/2 [22], we hypothesized that these residues are evolutionarily conserved due to their ability to modulate Scrt2 function through phosphorylation.

To test this hypothesis, we generated a series of single mutants at residues 77 and 78 to simulate the changes in residue charges prior and after phosphorylation. We replaced the original amino acids either with the neutral residues closest in structure to tyrosine or serine, or with acidic residues. Thus, Y77 was replaced with phenylalanine (*cScrt2*-Y77F) or glutamate (*cScrt-*Y77E) and S78 with alanine (*cScrt*-S78A) or aspartate (*cScrt2*-S78D). All four single mutations impaired Scrt2-mediated transcriptional repression (Fig. 3M).

**Figure 3.**
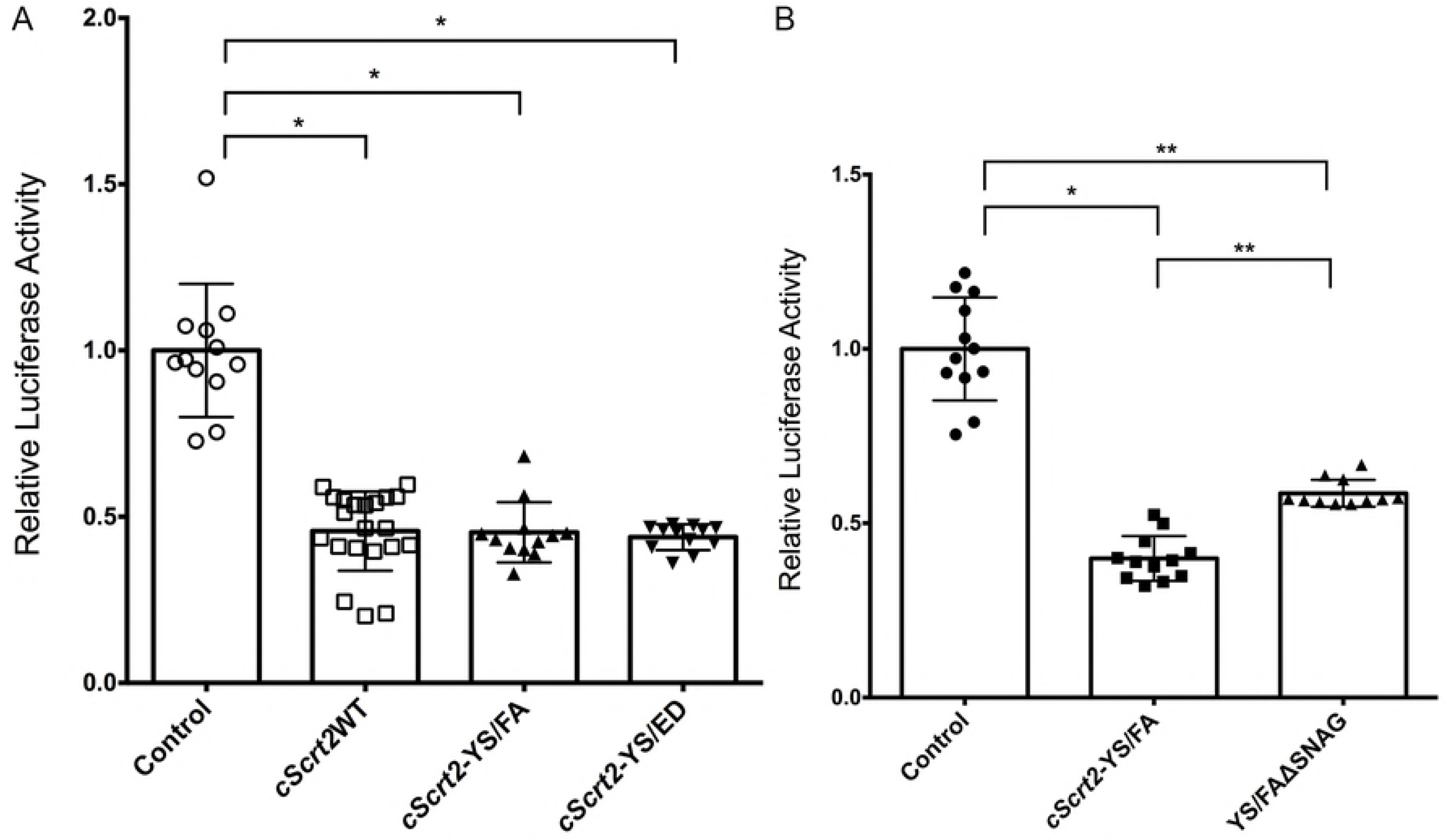
Mutations in residues Y77 or S78 do not alter the subcellular localization of Scrt2 but reduce the transcriptional repression activity. FLAG-tagged *cScrt2*-Y77F (A-C), *cScrt2*-Y77E (D-F), *cScrt2*-S78A (G-I) and *cScrt2*-S78D (J-L) remain in the nucleus of HEK293T cells after transfection. FLAG immunostaining co-localizes with nuclear GFP and DAPI. *cScrt2*-Y77F, *cScrt2*-Y77E, *cScrt2*-S78A were inserted in a pCIG vector whereas *cScrt2*-S78D was inserted in a pMES vector. (M) The single mutant forms *cScrt2*-Y77F, *cScrt2*-Y77E, *cScrt2*-S78A and *cScrt2*-S78D have reduced transcriptional repression activity compared to *cScrt2*WT (**), although the remaining activity is sufficient to produce a significant reduction in luciferase signal when compared to control (*). Results shown are the mean of 3 independent experiments performed on triplicate samples. The data were analyzed with one-way ANOVA non-parametric pairwise comparison and are represented as mean with standard deviation. * p<0.001; ** p=0.047; *** p=0.0164; @ p=0.04; # p= 0.006. Scale bar: 20μm

Reduction of transcriptional activity could not be attributed to changes in Scrt subcellular localization, as all constructs were found in the nucleus (Fig. 3B, E, H and K). Moreover, these mutations did not change protein expression levels (data not shown).

As our homology analysis suggested a co-evolution of the HINGE and SCRATCH domains, we hypothesized that the two domains act together, which would mean that removing the SCRATCH domain in the background of Y77 or S78 single mutants should further decrease Scrt-2 repressor activity. Contrary to our hypothesis, removal of the SCRATCH domain restored the repressor activity of *cScrt2*-Y77F and *cScrt*-S78A (Fig. 4).

**Figure 4.**
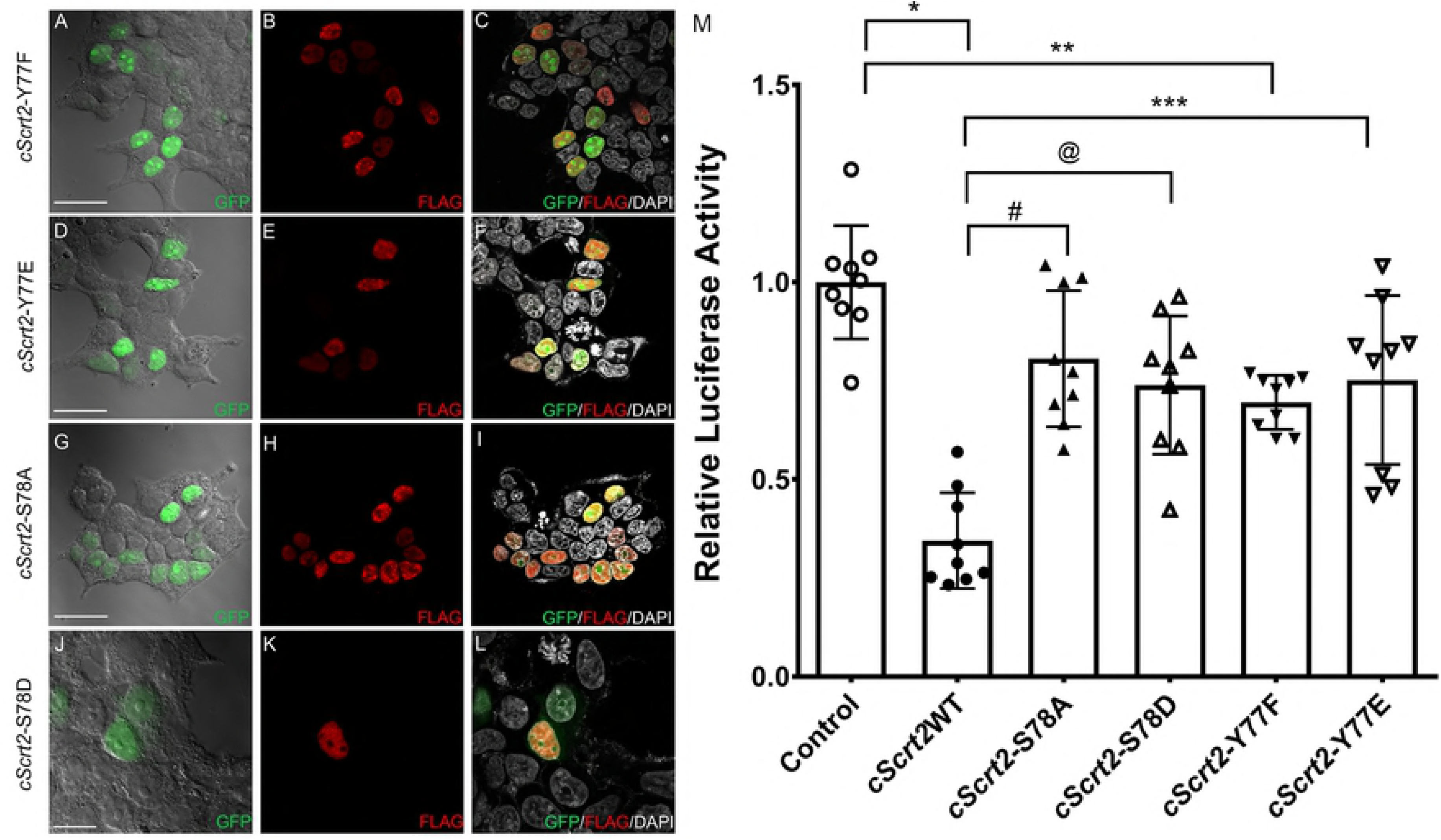
Mutation of Y77 or S78 does not reduce transcriptional repressor activity in the absence of SCRATCH domain. Mutation of the Y77 or of the S78 residues in the absence of the SCRATCH domain (YFΔSCRATCH or SAΔSCRATCH) does not alter the chicken Scrt2 transcriptional repression activity. Results shown are the mean of 5 independent experiments performed on triplicate samples. Statistical significance was calculated using one-way ANOVA multiple comparisons. *p<0.0001; **p=0.02; #p=0.0001; @p=0.0009.

### Double mutants of the HINGE domain repress transcription

Considering that single mutations of either Y77 or S78 decreased Scrt2 repressor activity and that invertebrates lack the entire HINGE domain (Fig. 2), we next tested the effect of simultaneously mutating both sites in Scrt2 (*cScrt2-*YS/FA and *cScrt2-*YS/ED). These double mutants did not differ significantly from wild type Scrt2 in their ability to repress transcription (Fig. 5A).

**Figure 5.**
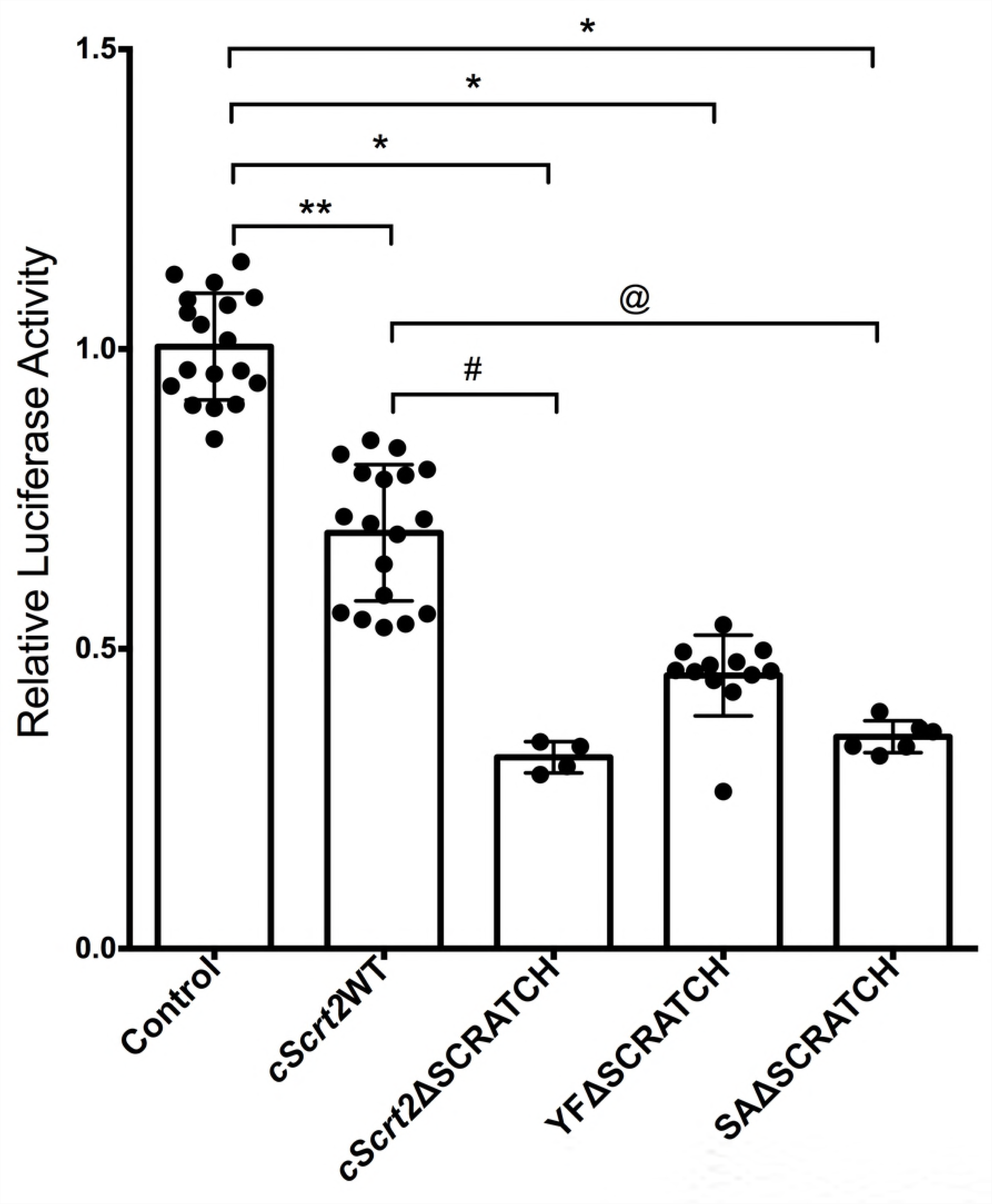
Simultaneous mutations in residues Y77 and S78 do not affect chicken Scrt2 activity. Transcriptional repression activity of the double mutant forms, *cScrt2*-YS/FA and *cScrt2*-YS/ED is similar to *cScrt2*WT (A). Removal of the SNAG domain (YS/FAΔSNAG) partially reduces the repressor activity of the double mutant form *cScrt2*-YS/FA (B). Results shown are the mean of 3 independent experiments, performed on triplicate samples. Statistical significance was calculated using 1-way ANOVA. *p<0.0001;**p=0.02

Considering the importance of the conserved SNAG domain in transcriptional repression, we next asked if the double mutants also repressed transcription through this domain. Indeed, when we compared the repressor activity of the double mutant (c*Scrt2*-YS/FA) in the presence or absence of the SNAG domain (ΔSNAG), the absence of SNAG partially decreased repressor activity in the double mutant (Fig. 5B).

## Discussion

We have dissected here the contributions of evolutionarily conserved domains in Scrt2 towards its transcriptional repression activity. In particular, we focused on the importance of the SNAG and SCRATCH domain and identified a novel conserved region called HINGE. Removal of SNAG and single-residue mutations in HINGE domain downregulated transcriptional repression.

Extensive phylogenetic comparison of the SNAG domain in the Snail superfamily suggests that it can be subdivided into two separate subdomains: SNAG-1 and SNAG-2. SNAG-1 is defined as the small domain of highly conserved residues, also known as the minimal SNAG (MPRSFLVKK), whereas SNAG-2 contains the subsequent 13-17 amino acids [21]. SNAG-1 and SNAG-2 do not necessarily occur in the same protein; in other words, the two subdomains can evolve independently, suggesting that they contribute to different functions. Scratch proteins - including Scrt2- all lack a recognizable SNAG-2 subdomain. However, Scrt2 retains a SNAG-1 subdomain identical to that found in Snail1. Our data shows that removal of SNAG-1 decreased Scrt2 repressor activity but not protein stability. In light of this, a more complex picture of SNAG-1 and 2 function arises. The canonical model of SNAG-mediated repression in the Snail superfamily relies on the functional analysis of Snail1, after simultaneously removing SNAG-1 and 2. In these experiments, repressor activity was completely abolished, and protein stability reduced [12,13]. Repressor activity has been attributed to the interaction of individual residues in SNAG-1 to repressor proteins and epigenetic modifiers. For example, Lysine-specific demethylase 1 (LSD1) interacts with Pro_2_ and Arg_3_ of Snail´s SNAG-1 domain [12,13]. Further, the Ajuba family of LIM repressor proteins interacts with Phe_4_ in SNAG-1 [23]. Together, these data suggest that the SNAG-1 domain is the minimal domain required for transcriptional repression, whereas SNAG-2 might be more relevant for Snail1 protein stability.

We also identified and analyzed the role of another evolutionarily conserved domain (HINGE), that lies between the SNAG and SCRATCH domains, containing potential phosphorylatable sites. As phosphorylation of Snail1 and 2 modulate their stability and repressor activity [22], we explored the importance of these residues for Scrt2 function through point mutations. Our mutations focused in changing the charge of the original residues, substituting them by the neutral residues alanine and phenylalanine or by the negatively charged residues aspartate and glutamate: substitution with alanine or phenylalanine generates a non-phosphorylatable form, whereas substitution with aspartate or glutamate simulates the negative charge provided by phosphorylation [22,24]. Thus, if phosphorylation at these residues modulated Scrt2 activity, negatively charged and neutral charge point mutations should yield opposite results. Instead, all single point mutations decreased Scrt2 transcriptional repressor activity irrespective of the residue charge, suggesting that the replacement of these amino acids affects Scrt2 activity through changes in protein conformation. Without definitive crystallographic information, our interpretation of the single and double mutant data is that this region acts as a hinge. Interestingly, double mutants restored repressor activity, but in a SNAG-dependent manner: with SNAG deleted, double mutations did not restore transcriptional repression. Thus, the double mutations might have rearranged protein conformation so as to expose SNAG in a position that allows co-repressor recruitment.

The mutations in the HINGE domain also revealed a putative modulatory role for the SCRATCH domain on Scrt2 function: in single mutations of either residue, 77 or 78 of HINGE, concomitant removal of the SCRATCH domain restored transcriptional repressive activity. However, removal of SCRATCH did not affect Scrt2 activity in the background of an intact HINGE domain containing Tyrosine77 and Serine78. Thus, the modulatory activity of SCRATCH depends on the identity of the residue on position 77 or 78 at the HINGE domain, suggesting that their function evolved in concert. Indeed, our phylogenetic analysis indicate that the HINGE domain co-evolved with the SCRATCH domain. Conservation of the SCRATCH domain is higher amongst species that contain both Tyrosine and a Serine in the HINGE domain (Fig. S2). If true, the salmon and zebrafish Scrt2 orthologues, which present the Tyrosine but not the Serine residue in the HINGE domain (EEYCD), might present a conformation where SCRATCH is constitutively modulating transcriptional repression, and thus would present a lower activity than their avian or mammalian counterparts. In the case of these latter species, which have fully conserved HINGE domains, post-translational modifications in HINGE could change its conformation so as to activate the modulatory role of SCRATCH. In this scenario, addition of a single negative charge, possibly through phosphorylation at position 77 or 78, would be sufficient to activate SCRATCH-domain-mediated reduction of Scrt2 transcriptional repression. Although our data shows that changing residue 77 or 78 to a neutral aminoacid has the same effect as a substitution for a negatively-charged one, we cannot rule out the possibility that experimental substitutions of aminoacid residues fail to completely reproduce the changes triggered by phosphorylation.

Finally, we also investigated the role of the zinc-finger domain in nuclear translocation. The chicken Scrt2 zinc-finger domain has 61.47% identity to the homologous region in mouse Snail1 and was sufficient to promote nuclear localizationScrt2. The nuclear shuttling function of mouse Snail1 is attributed to importin binding to six basic and six hydrophobic residues [25]. Although the zinc-finger domain in Scrt2 contains all the six importin-binding hydrophobic residues, it lacks one of the importin-binding basic residues identified in Snail1 (Fig. S4), indicating that conservation of five of the basic residues is sufficient for nuclear localization. Also, the zinc-finger domain is sufficient to direct protein-DNA interaction at E-box motifs (Fig. S5).

Thus, we confirm that Scrt2, with a general structure similar to the Snail family members, relies on SNAG for transcriptional repression and the zinc-finger domain for nuclear translocation and DNA-binding. We also show that Scrt2 has additional domains that modulate transcriptional repression. Together, our data extends current knowledge on the modular structure of Snail superfamily members and provides support for the hypothesis that modularity in this superfamily arose from duplication and divergence from a common ancestral protein.

## Acknowledgements

The authors thank Dr. Paulo Sérgio Lopes Oliveira for discussions about protein structure, Dr. Cristóvão Albuquerque for editing assistance and Dr. Ali Brivanlou for generously sharing lab space and reagents.

## Supporting information

Supplementary figure 1

**Chicken Scrt2 localizes to the nucleus in chick neural tube cells**. Immunostaining of neural tube sections show presence of MYC or FLAG tags in the nucleus 24 hours after electroporation with MYC-tagged *cScrt2*WT (B) or FLAG-tagged *cScrt2*-Y77E (E) and *cScrt2*-S78A (H). MYC (B) and FLAG (E-H) signal co-localizes with GFP (A, D, G) and DAPI (C, F, I). C’, F’ and I’ are a higher magnification of the overlap image in the dotted area in B, E and H. Vector reporter GFP labels the electroporated cells. Scale bar – 50 μm.

Supplementary figure 2

**HINGE and SCRATCH domains co-evolved in vertebrates and were both modified in fish.** (A) The HINGE-SCRATCH region of different species was aligned to compare the changes in both domains simultaneously. Conservation of the SCRATCH domain is higher amongst species that contain both Tyrosine and a Serine in the HINGE domain. (B) Full length SCRT2 amino acid sequences were aligned in CLUSTALX and the resulting N-J tree rooted with the wasp SCRT2 sequence. Python and Baiji sequences were partial and lacked the SNAG domain.

Supplementary figure 3

**Scrt2 has potentially phosphorylated residues.** *In silico* analysis of the chicken Scrt2 sequence using the online phosphorylation prediction site KinasePhos identified residues Y77, S78 and S82 as possible targets for phosphorylation. Below are candidate kinases for these phosphorylation sites. The box outlined in dashed light blue lines shows the Scrt2-specific domain (aa 75-85 in chicken Scrt2).

Supplementary figure 4

**Alignment of the zinc-finger domains of selected members of the Snail superfamily.** ClustalW alignment of the region containing zinc-fingers 2-4 of Snail and Scratch orthologues is shown here. The residues that interact with importin are highlighted with different colors: basic residues are red and hydrophobic residues are green. The labels are preceded by the species; h: human, m: mouse, c: chicken, d: Drosophila. Sequences used were hSnail1 (NP_005976.2), hSnail2 (NP_003059.1), hScratch1 (NP_112599.2), hScrt2 (NP_149120.1), mSnail1 (NP_035557.1), mSnail2 (NP_035545.1), mScratch1 (NP_570963.1), mScrt2 (NP_001153882.1), cSnail1 (NP_990473.1), cSnail2 (CAA54679.1), cScrt2 (AEW43643.1), dSnail (NP_476732.1), dScratch (AAD38602.1).

Supplementary figure 5

**Chicken Sctr2 represses transcription driven by E-box.** HEK293T cells were transfected with pGL3- 4xE-box and empty pMES plasmid (Control), or full length Scrt2 (*cScrt2*WT) or Scrt2 zinc-fingers fused to the repressor domain of Engrailed (EN-*Scrt2*) or to the VP16 activator domain (VP16-*Scrt2*). VP16- *Scrt2* strongly enhanced transcriptional activity (t-test, p<0.001) while *cScrt2*WT reduced transcription below the basal levels (t-test, p<0.05); EN-Scrt2-mediated reduction was not significantly different from *cScrt2*WT.

